# Targeted and selective knockout of the TLQP-21 neuropeptide unmasks its unique role in energy homeostasis

**DOI:** 10.1101/2023.03.23.532619

**Authors:** Bhavani S. Sahu, Maria Razzoli, Seth McGonigle, Jean Pierre Pallais, Megin E. Nguyen, Masato Sadahiro, Cheng Jiang, Wei-Jye Lin, Kevin A. Kelley, Pedro Rodriguez, Rachel Mansk, Cheryl Cero, Giada Caviola, Paola Palanza, Loredana Rao, Megan Beetch, Emilyn Alejandro, Yuk Y. Sham, Andrea Frontini, Stephen R. Salton, Alessandro Bartolomucci

## Abstract

Pro-peptide precursors are processed into biologically active peptide hormones or neurotransmitters, each playing an essential role in physiology and disease. Genetic loss of function of a pro-peptide precursor results in the simultaneous ablation of all biologically-active peptides within that precursor, often leading to a composite phenotype that can be difficult to align with the loss of specific peptide components. Due to this biological constraint and technical limitations, mice carrying the selective ablation of individual peptides encoded by pro-peptide precursor genes, while leaving the other peptides unaffected, have remained largely unaddressed. Here, we developed and characterized a mouse model carrying the selective knockout of the TLQP-21 neuropeptide (ΔTLQP-21) encoded by the *Vgf* gene. To achieve this goal, we used a knowledge-based approach by mutating a codon in the *Vgf* sequence leading to the substitution of the C-terminal Arginine of TLQP-21, which is the pharmacophore as well as an essential cleavage site from its precursor, into Alanine (R_21_→A). We provide several independent validations of this mouse, including a novel in-gel digestion targeted mass spectrometry identification of the unnatural mutant sequence, exclusive to the mutant mouse. ΔTLQP-21 mice do not manifest gross behavioral and metabolic abnormalities and reproduce well, yet they have a unique metabolic phenotype characterized by a temperature-dependent resistance to diet-induced obesity and activation of the brown adipose tissue.

## Introduction

Genetic loss of function is a powerful approach to identify the necessary role of gene products on cellular functions, physiology or diseases. While this approach is common for genes encoding a single protein product (receptors, structural proteins, signaling proteins etc.), pro-peptide precursor genes pose a serious challenge because they encode multiple biologically-active peptides each having a different function and unique signaling ^1–4^. Indeed, all of the encoded peptides are simultaneously ablated using standard knockout approaches or cre-lox systems, resulting in a mixed phenotype which is difficult to explain from the loss of individual parts. Propeptides precursors such as pro-opio-melanocortin (POMC), pro-ghrelin, pro-glucagon, brain-derived neurotrophic factor, several granin peptides and many others, are key regulators of neuroendocrine functions and play essential roles in physiology and disease ^1–4^. Notably, these peptides can be active on different receptors and exert very different biological functions. Classic examples include adrenocorticotropic hormone (ACTH), α–Melanocyte-stimulating hormone (αMSH) and β-endorphin, which are all derived by differential processing of the same POMC precursor ^3^. Additionally, their tissue processing it tightly regulated by proteases leading to tissue-specific prevalence of only one or a few of those peptides. Lastly, to further complicate the biology of this system, some of these pro-peptide precursors also play a critical role in dense core granule biogenesis and stability ^1,5,6^. Due to these biological constraints and technical limitations, mice carrying the selective ablation of only one peptide, encoded by a pro-peptide precursor gene leaving the other peptides unaffected remains largely unaddressed. For example, while a knockout of the POMC precursor was developed long ago ^7^, there is no selective ACTH or β-endorphin knockout that would also express the other unaltered POMC-derived peptides. One exception is a point mutant mouse in which a KKRR→QKQR mutation in the cleavage site of ACTH prevents the production of α-MSH ^8^. In general, alternative approaches have been preferred, including the use of selective immune-neutralizing antibodies ^2^, and peptide injection into propeptide knockout mice ^3^. These alternative approaches have limitations including specificity and bioavailability of the antibodies, and pharmacological properties of the drugs injected into knockout mice. Standard gene targeting approaches, CRISPR-Cas9, and other recent gene editing technologies, are all potentially useful approaches to address this limitation, each with different advantages and disadvantages ^9^.

The *Vgf* gene (non-acronymic) encodes for a 615 amino acid (617 in rodents) long pro-protein precursor of at least 8 biologically-active peptides ^1^. VGF is cleaved by prohormone convertases (PC)1/3 and PC2 to generate low molecular weight peptides which are secreted through the regulated pathway ^1,10^. Thus - much like for POMC - germline ^11,12^, or tissue-specific ^6,13,14^ VGF null mice, or humanized knock in mice in which the human *VGF* coding sequence replaces the mouse coding sequence in the mouse *Vgf* gene locus ^15^, are not informative about the biology of any individual VGF-encoded peptides. Additionally, the physiology of VGF is also complicated by the structural role of the pro peptide which is likely relevant for dense-core granule biogenesis and secretion of other peptides by endocrine cells (e.g. insulin in beta-cells) ^1,6^. Among the VGF-derived peptides, the C-terminal internal 21 amino acid long fragment known as TLQP-21 ^16^, is arguably the most studied and best characterized; also being the only one for which a clear receptor-mediated mechanism has been identified thus far ^17–19^. First identified in the rodent brain ^16^, and later in sympathetic and sensory nerves ^20,21^ and in endocrine glands such as the adrenal medulla ^22^, TLQP-21 emerged as a pleiotropic neuropeptide involved in various physiological processes such as lipolysis, microglial activation, pain, sexual behavior and energy balance ^1,17^. However, most of our knowledge on TLQP-21 derives from pharmacological gain of functions experiments, while selective loss of function approaches have not been attempted yet. Here, after reconstructing the evolutionary history of TLQP-21 highlighting a strong selective pressure on its pharmacophore in mammals, we generated and initially characterized a TLQP-21-selective knockout mouse (aka, ΔTLQP-21). We took advantage of previous knowledge demonstrating that the C-terminal Arginine, which is necessary for TLQP-21 biological activity ^17,19,23^, is also required for cleavage from its precursor TLQP-62 ^10,20,24^. We mutated the codon in the *Vgf* sequence encoding for the C-terminal Arginine into a codon encoding Alanine (R_21_→A) which results in the production of a peptide sequence that is not a substrate for pro-hormone convertase ^25^. This resulted in a mouse lacking TLQP-21 while showing normal expression of pro-VGF and the TLQP-21 precursor TLQP-62. ΔTLQP-21 mice do not have gross behavioral and metabolic abnormalities, yet they manifest an unexpected metabolic phenotype characterized by a temperature-dependent resistance to diet-induced obesity.

## Results

### The evolutionary history of TLQP-21 neuropeptide suggests that an intense selective pressure occurred at its C-terminus in mammals

Recent evidence suggests that C3a/TLQP-21/C3aR1 genes underwent intense selective pressure in mammals ^23^. Specifically, while C3a was identified as the ancestral ligand of C3aR1 based on sequence analysis and is highly conserved across mammals ^23^, TLQP-21 pharmacophore and C3aR1 binding pocket were subject to mutation during evolution which conferred a gain of function in the *Murinae* subfamily of rodents ^23^.

We have now expanded our evolutionary analysis on TLQP-21 (and also included C3a and C3aR1) including 161 mammalian sequences found in NCBI and focused on the C-terminal amino acid motif Proline-Proline-Alanine-Arginine (−PPAR_21_), the essential amino acids required for receptor binding and activation ^19^, with the R_21_ also representing a recognition site for prohormone convertase to cleave TLQP-21 from its precursor TLQP-62 ^1^ and for putative plasma proteases to deactivate the peptide ^1,19,23,24^. We found that R_21_ is highly conserved (68% of species) and that the amino acid Serine (S) prevails at position 20 in mammals including humans, and can thus be considered the ancestral sequence (^23^ and **Supplementary Table 1**). Conversely, A_20_ is unique to *Murinae/Cricetinae* (**Figure 1 A,B** and **Supplementary Table 1**). Interestingly, there are only 2 exceptions to the S/A_20_R_21_ motif (**Figure 1C**): i) marine mammal species (Artiodactyla: Balaenopteridae, Delphinidae, Monodontidae, Phocoenidae, Physeteridae) invariably express S_21_; ii) 38 species including carnivores (Canidae, Felix, Mustelidae), ursidae and horses express histidine (H) at position 21 (**Supplementary Table 1**). Neither H or S are canonical consensus sites for pro-hormone convertase ^25^, thus suggesting that TLQP-21 might not be produced in these species. This evolutionary sequence analysis indicates that selective pressures acted in different species on amino acids critical for TLQP-21/C3aR1 receptor recognition, while the C-terminus of C3a remains largely conserved across mammals (**Figure 1**). Yet, the functional significance of those mutations remains largely unknown. Here we aimed to shed light on this question by generating a mouse model in which TLQP-21 cannot be cleaved from its precursor TLQP-62.

**Figure 1.**
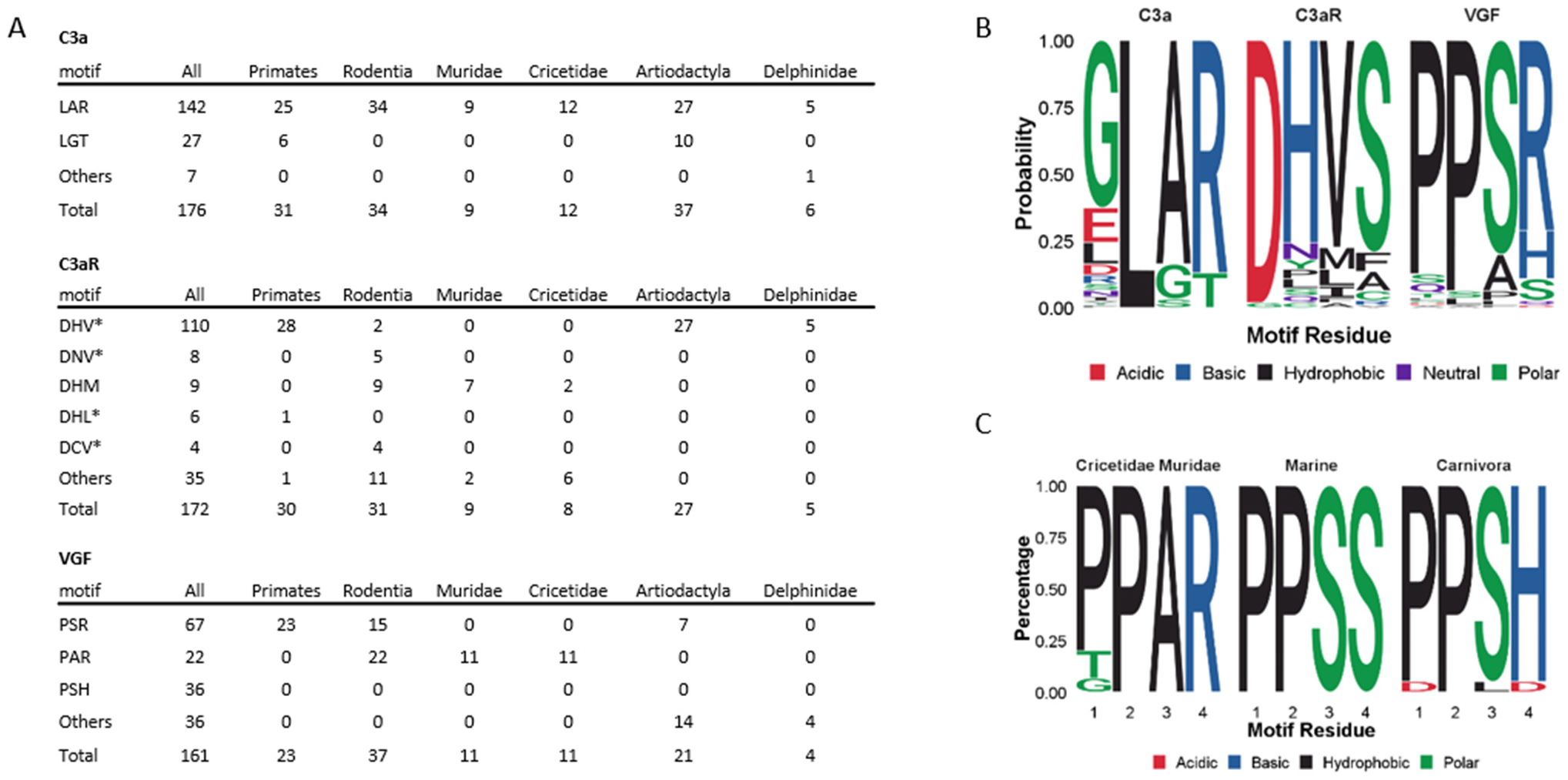
Evolutionary analysis of C-terminal TLQP-21 sequence. **A)** Table of C-terminal C3a and TLQP-21 and selected residues in the C3aR1 binding pocket motifs from NCBI Protein RefSeq database. **B)** Sequence logos of mammals for C3a C terminus, C3aR1 binding pocket and TLQP-21 pharmacophore. **C)** Sequence logos of mammals for the last 4 amino acids of TLQP-21 in selected groups of mammals. See Supplementary Table 1 for a complete sequence list.

### Generation and characterization of ΔTLQP-21 mice

TLQP-21 is a C-terminal internal fragment, which can only be produced upon cleavage of the precursor TLQP-62 ^16^. By utilizing gene targeting constructs previously employed to generate the VGF knockout mice ^11^ and inserting flanking loxP sites into the mouse Vgf locus ^26^, we introduced a mutation in the VGF coding sequence (P_573_PAR_576_ ccacctgcgcgc to P_573_PAA_576_ ccacctgcggcg) leading to R_21_→A mutation (**Figure 2A**). Sanger sequencing confirmed the correct mutation in the ΔTLQP-21 mutated mouse (**Figure 2B**). Staining the adrenal gland of WT and ΔTLQP-21 mice with a selective rabbit-anti-mouse TLQP-21 antibody ^20^ confirmed the loss of TLQP-21 in ΔTLQP-21 animals (**Figure 2C**). This mutation is predicted to result in the inability of pro-hormone convertases to cleave the peptide from TLQP-62 ^16^. To verify that this mutation did not impact the pro-peptide processing in general, we blotted brain extracts using a C-terminal antibody which recognizes the VGF pro-peptide and all precursor fragments containing the C-terminus, including TLQP-62 ^27^. Results show an identical profile of VGF fragments in WT and ΔTLQP-21 mice (**Figure 2D**). This result suggests that preventing the cleavage of TLQP-21 does not impact the pro-peptide nor does it increase the accumulation of its precursor TLQP-62. Lastly, expression of *Vgf, C3aR1* and *C3* in several key organs was unaffected by the ΔTLQP-21 mutation (**Supplementary Figure 1**). To confirm the R_21_→A Δ mutation at the protein level we developed a novel in-gel digestion-based MS analysis for TLQP-21 using an approach similar to what was used for other peptides ^28^ (**Figure 3A**). Fresh adrenal gland was rapidly lysed in RIPA lysis buffer and equal amounts of proteins were loaded in a 12% precast gel and were resolved in the MES buffer system along with the synthetic TLQP-21 peptide. Gels were stained with Safe blue followed by excision of bands between ∼2KD to ∼5KD corresponding to a region where TLQP-21 would be identified based on its molecular weight (MW) ^16^. Excised bands followed tryptic digestion and were analyzed by targeted mass spectrometry (MS). The point mutation R_21_→A in the VGF coding sequence results in the loss of a prohormone processing site leading to new peptide fragment transitions predicted by the Skylab software, such as AQAR and the unnatural HFHHALPPA<u>A</u>HHPDLEAQAR (**Figure 3B**). Our MS analysis reveals that those fragments can only be detected in the ΔTLQP-21 mice while being absent in the WT (**Figure 3C,D**; **Supplementary Figure 2**), overall validating the mouse model at the protein level.

**Figure 2.**
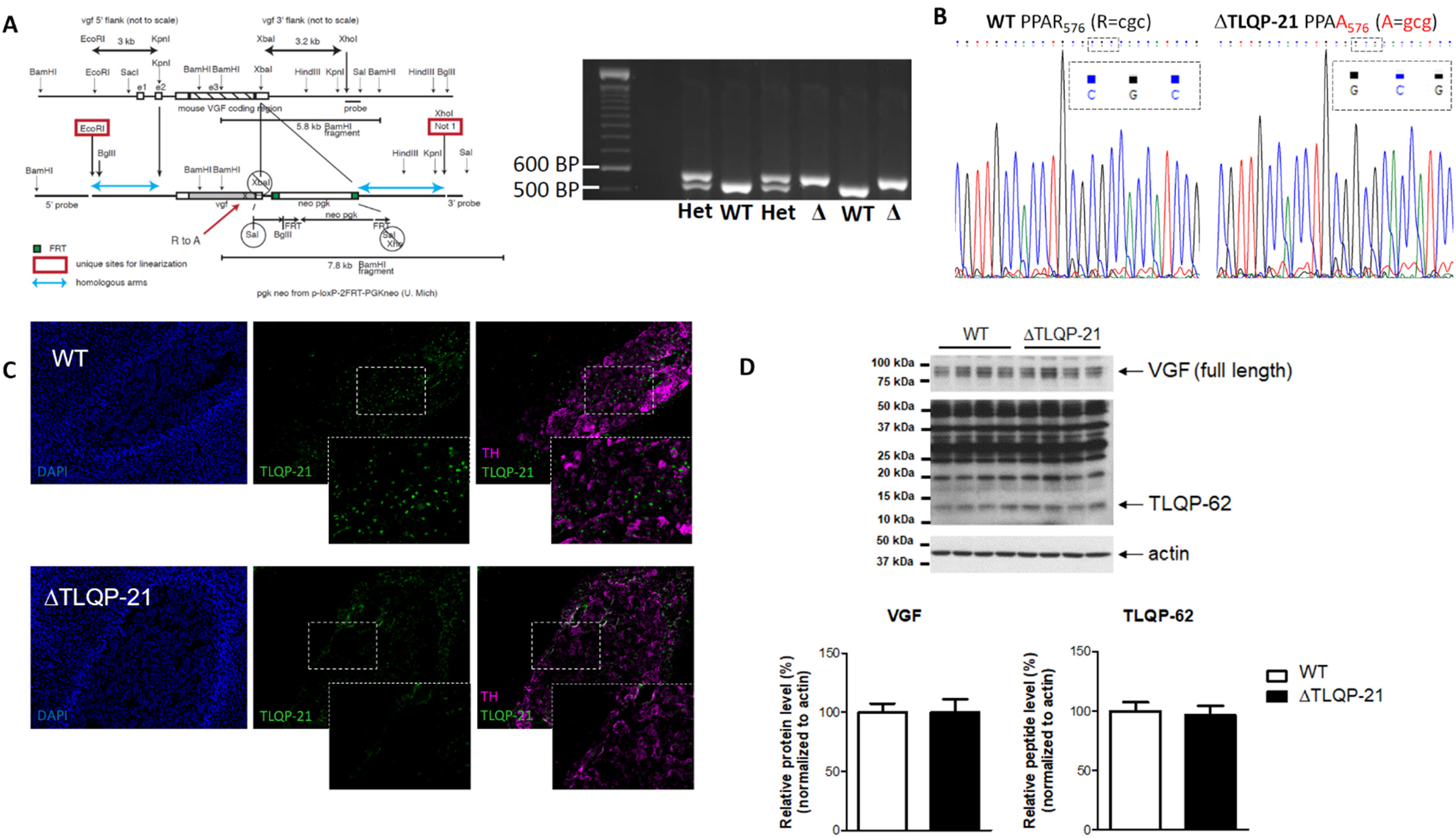
Generation and characterization of ΔTLQP-21 mice. **A)** Schematic for the generation of delta TLQP-21 mice and PCR amplification. Specific primers were designed to amplify the genomic delta TLQP-21 region of the *Vgf* gene, followed by gel extraction of the purified product, which reveals a single specific ∼180 base pair amplicon of this VGF region. **B)** Sanger sequencing. The TLQP-21 PCR amplified region was gel purified and subjected to Sanger sequencing. The results were analyzed using Chromas Lite software, and base changes in the delta mice were confirmed by manually inspecting the peaks of the chromatogram. **C)**. IHC of mouse adrenal gland. Sections of the adrenal gland were stained with antibodies for TLQP-21 (rabbit anti-TLQP-21, 1:100, ^20^) or Tyrosine Hydroxylase (sheep polyclonal TH, 1:100, Millipore Sigma). Images representative of N=3. **D)**. Western blotting of the hypothalamus. A VGF C-terminal specific primary antibody was used to detect on the same blot with different exposure lengths, the VGF pro-protein (top panel, 75-100 kDa), and the TLQP-21 precursor TLQP-62 (middle panel, 10-15 kDa), along with a number of additional C-terminal-containing VGF-derived peptides (middle panel, 15-50 kDa). The actin loading control was visualized in the lower panel (37-50 kDa). Relative levels of VGF and TLQP-62 normalized to actin, for WT and ΔTLQP-21 samples (WT = 100%) were analyzed by Student’s t-test (ns, N=4).

**Figure 3.**
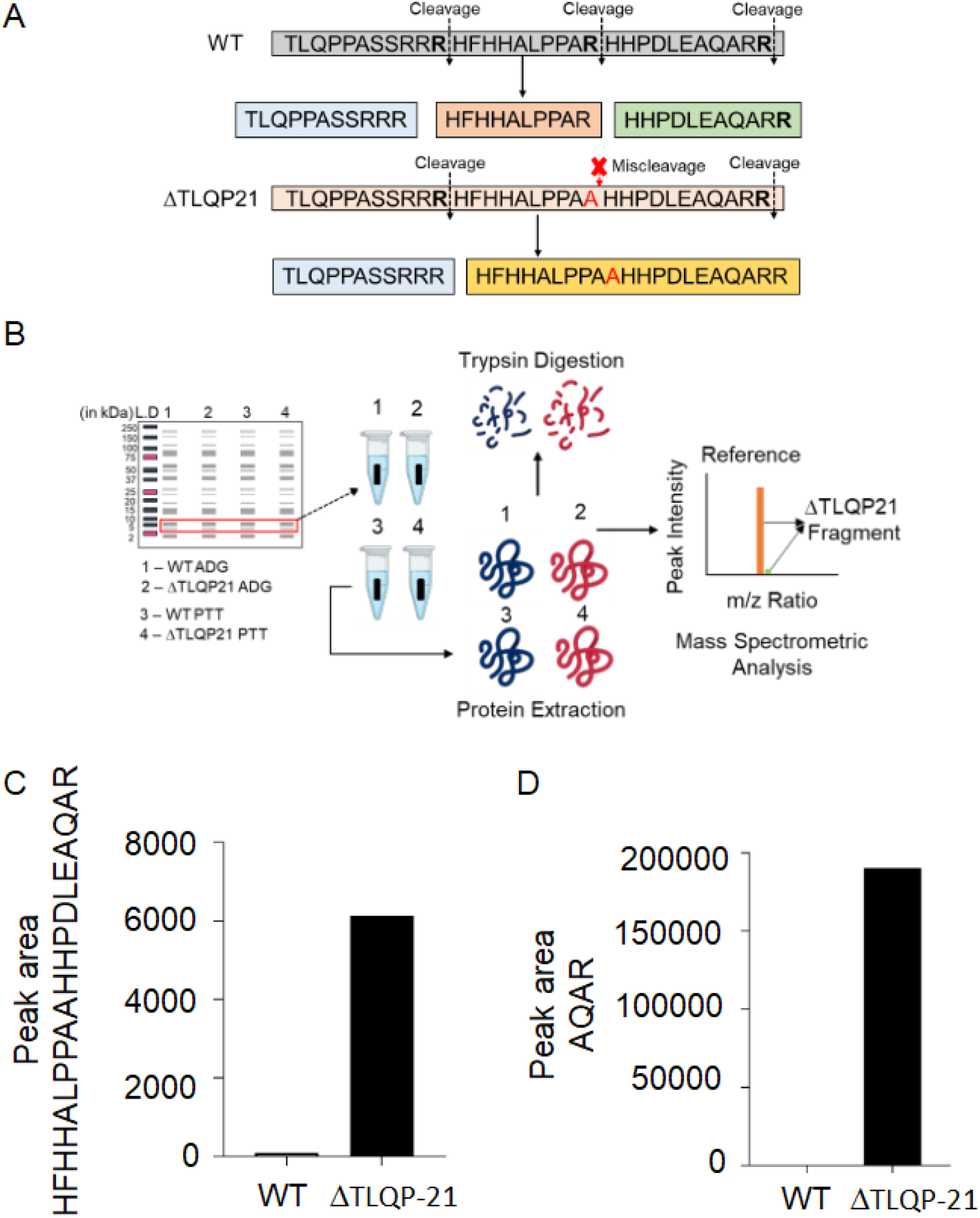
Targeted mass spectrometry. **A)**. A targeted mass spectrometry-based approach was used to characterize the misprocessing of TLQP-21 in ΔTLQP-21 mice. Tissue samples were subjected to gel electrophoresis along with the synthetic TLQP-21 peptide followed by excision, tryptic digestion, and analysis by targeted mass spectrometry. **B)**. Schematic of WT and ΔTLQP-21 sequences showing predicted cleavage sites and the point mutation R→A. **C**,**D)**. Targeted mass spectrometry of the adrenal gland. The R→A mutation in the ΔTLQP-21 mice ablated a proteolytic processing site which resulted in a larger tryptic fragment HFHHALPPAAHHPDLEAQAR and the presence of AQAR which is normally not present in TLQP-21. Representative MS spectra of WT and ΔTLQP-21 are shown in Supplementary Figure 2.

### ΔTLQP-21 mice show normal development and behavior

TLQP-21 has been implicated in multiple biological functions^1,16,17,29,30^, thus we conducted a thorough characterization of ΔTLQP-21 mouse behavior and physiology. ΔTLQP-21 mice are viable, and homozygous or heterozygous mice can reproduce with a similar success rates and manifest similar maternal behavior (**Supplementary Tables 2-3**). Male and female ΔTLQP-21 mice from homozygous or heterozygous breeding pairs reached comparable weight at weaning suggesting similar developmental rates (**Supplementary Table 4**). Due to the known effects of pharmacological treatment with TLQP-21 on behavior ^14,31,32^, mice were also tested in traditional behavioral assays to detect anxiety-like and depression-like traits, but again, no differences were found compared to WT (**Supplementary Figure 3**). Based on a similar phenotype we opted to continue the experiments using pups from heterozygous pairs, allowing for better control of the early maternal environment, and using littermates for all the experiments.

### ΔTLQP-21 mice are resistant to diet-induced obesity at room temperature

One of the best-characterized functions of TLQP-21 is to potentiate adrenergic-induced lipolysis ^20,23,33^. Consistently, plasma free glycerol measured in response to 3h cold challenge at 4°C was decreased in ΔTLQP-21 male and female mice (**Supplementary Figure 4A**). Conversely, free glycerol induced by a β-adrenergic receptor (β3AR) agonist, which does not activate sympathetic transmission and thus TLQP-21 secretion from neurons, was unaffected by TLQP-21 deletion (**Supplementary Figure 4B**).

Based on the catabolic role exerted by TLQP-21 upon central or peripheral administration ^16,33,34^, we initially hypothesized that its selective deletion might result in obesity. At two months of age, male and female ΔTLQP-21 and WT mice were singly housed and monitored for their body weight growth and food intake in response to standard diet. No differences emerged for body weight, food intake, body composition, energy expenditure, insulin tolerance, glucose tolerance, and fasting glucose due to the mouse genotype (**Supplementary Figure 5 and 6**). Next, we challenged the ΔTLQP-21 mice with an obesogenic high fat diet (HFD). Contrary to our hypothesis, and similar to VGF-/- and C3aR1-/-mice ^11,35^, both male and female ΔTLQP-21 mice were resistant to diet induced obesity as demonstrated by significantly lower body weight gain and fat mass gain (**Figure 4 A-D**) without showing a distinct food intake, or fat free mass gain compared to WT (**Supplementary Figure 6**). Consistent with the leaner phenotype, plasma insulin was significantly lower, and insulin sensitivity was higher in ΔTLQP-21 mice than WT (**Figure 4E-H**). Unexpectedly, resistance to obesity was not paralleled by improved glucose tolerance or fasting glucose levels (**Supplementary Figure 6**).

**Figure 4.**
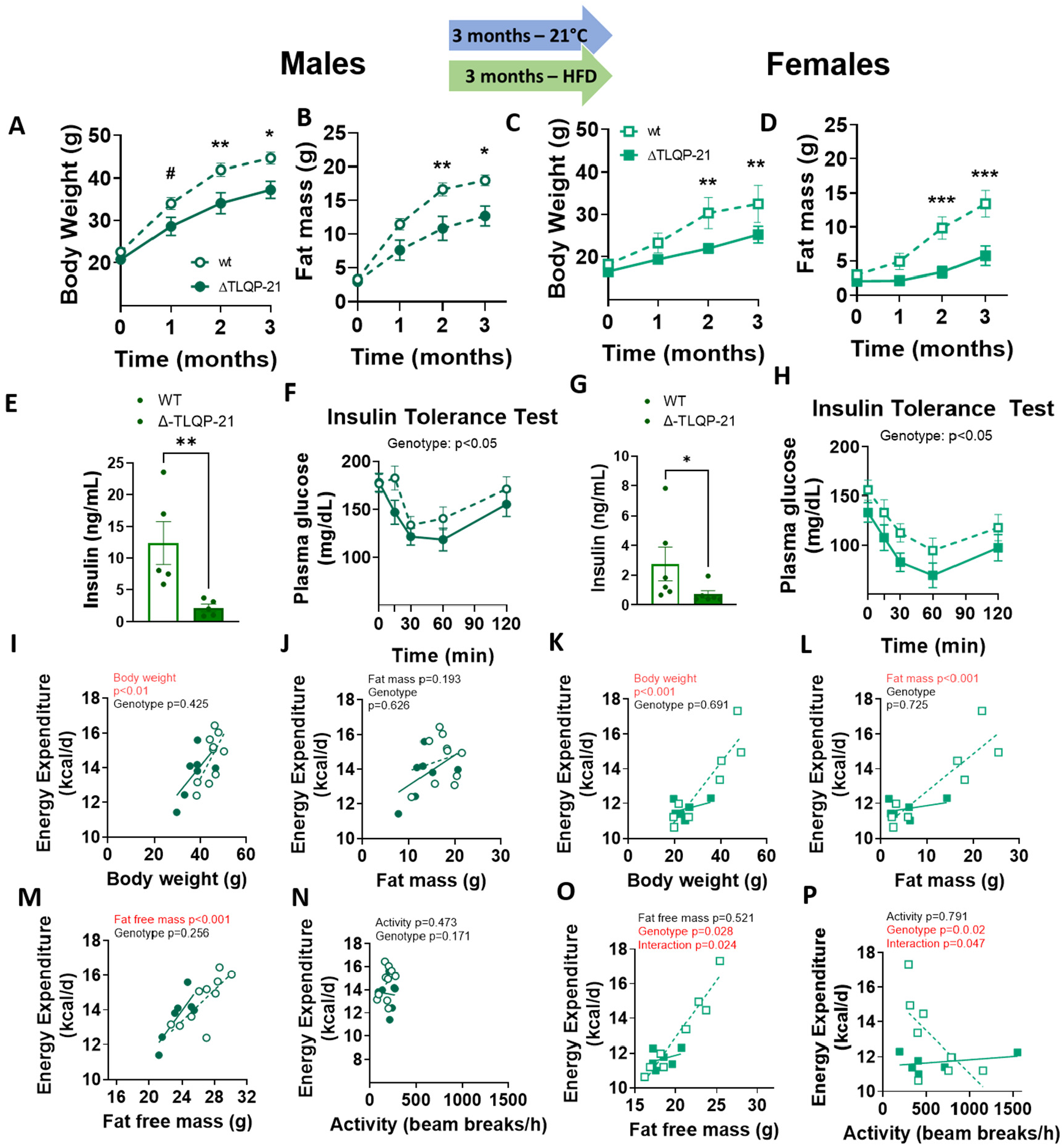
Metabolic phenotypes of ΔT21 mutant male and female mice and their wild type counterparts tested at room temperature (21±2C°) and ad libitum fed 60% High Fat Diet (HFD). Both male and female ΔTLQP-21 mice gained significantly less body weight (main effect genotype: F(1,32)=11.12, p<0.01; main effect sex: F(1,32)=39.86, p<0.0001; main effect time on diet: F(3,96)= 139.74, p<0.0001) as well as fat mass (main effect genotype: F(1,32)=13.75, p<0.001; main effect sex: F(1,32)=20.00, p<0.0001; main effect time on diet: F(3,96)= 85.35, p<0.0001) over time than WT mice (A-B,C-D). ΔTLQP-21 mutant mice of both genders showed a significantly lower non fasting insulin (Male: U=0; Female: U=4; note that data were not normally distribute d and were analyzed with the non-parametric Mann Whitney U Test) and greater insulin sensitivity (main effect genotype: F(1,34)=5.04, p<0.05; main effect sex: F(1,34)=20.71, p<0.0001) than WT animals (F,H). Energy expenditure over 24 h was significantly influenced by body mass in both males and females (I,K), by fat mass in females (L) and by fat free mass in males (M). In female mice, genotype significantly influenced energy expenditure that was lower in ΔTLQP-21 mice, while interacting with both fat free mass (O) and activity (P). N=6-9/group. Data in A-D and F,H were analyzed with repeated measure ANOVA with time point and genotype as main factors; symbols indicate differences between genotypes at each time point. #represents 1>p>0.05; * represents p<0.05; ** represents p<0.01. Data in I-P were analyzed using ANCOVAs with the factor indicated on the x axis as continuous predictor.

The brown adipose tissue (BAT) of the ΔTLQP-21 male mice fed HFD appeared markedly more activated, with brown adipocytes having smaller lipid droplets and higher level of UCP1 compared to WT (**Figure 5A,B**). Consistent with the morphological analysis, the BAT of male ΔTLQP-21 mice fed HFD showed increased *Ucp1, Cox71a, Zic1, and Adrb3* expression, while no differences emerged in mice fed a standard diet (STD) (**Figure 5C**). Morphologically, the perigonadal white adipose tissue (pWAT) but not the subcutaneous inguinal (sc)WAT of male mice fed HFD visually showed a higher proportion of smaller-size adipocytes in the ΔTLQP-21 mice, which is consistent with obesity resistance (**Supplementary Figure 6**). Fat pad morphology and gene expression of female ΔTLQP-21 was substantially similar to WT mice, with a prominent effect of HFD. The BAT of female mice had a more activated appearance than in males with smaller adipocytes and higher levels of UCP1, and effect which was more evident in ΔTLQP-21 mice (**Figure 5; Supplementary Figure 6**).

**Figure 5.**
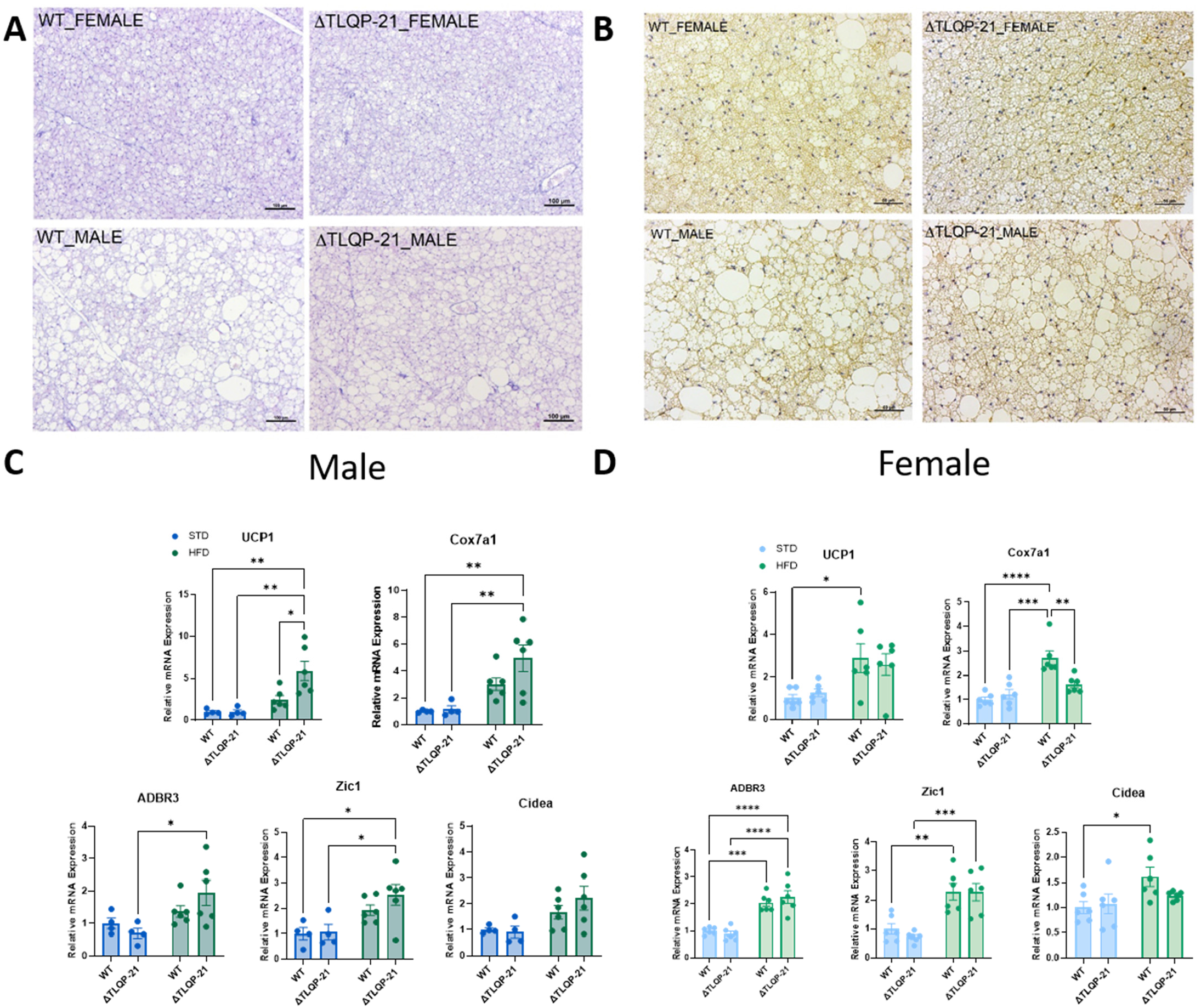
Brown adipose tissue morphological and molecular analysis. **A)** H and E staining and **B)** IHC for UCP1 in BAT obtained from male (bottom row) and female (top row) in WT and ΔTLQP-21 mice fed with HFD. **C**,**D**). QPCR of genes implicated in brown adipocyte thermogenesis and browning in WT and ΔTLQP-21 male and female fed standard diet (STD) or a high fat diet (HFD) and housed at room temperature. UCP1 (Males: genotype F(1,16)=4.737, p=0.0448, diet F(1,16)=16.34, p=0.0009, interaction F(1,16)=4.764, p=0.0443; Female: diet F(1,20)=13.14, p=0.0017), Cox7a1 (Male: genotype F(1,20)=4.892, p=0.0388, diet F(1,20)=30.05, p<0.0001; interaction F(1,20)=11.46, p=0.0029; Females: diet: F(1,16)=17.76, p=0.0007), ADRB3 (Males: diet F(1,16)=8.298, p=0.0109; females: diet F(1,20)=69.05, p<0.0001), Zic1 (Males: diet F(1,16)=13.48, p=0.0021; females: diet: F(1,20)=37.61, p<0.0001), Cidea (Males diet: F(1,16)=8.387, p=0.0105; females: diet: F(1,20)=6.580, p=0.0185). Data in C,D were analyzed with two-way ANOVA with diet and genotype as main factors with Tukey’s multiple comparisons test as ad hoc analysis. Symbols represent individual ΔΔCt values normalized over WT at STD. * represents p<0.05, ** represents p<0.01, *** represents p<0.001, and **** represents p<0.0001.

The resistance to HFD-induced obesity and the BAT morphology in ΔTLQP-21 mice, suggest increased energy expenditure. However, heat production measured via indirect calorimetry at the end of the study was not increased in either males or females (**Figure 4 I-P**).

### The obesity resistance of ΔTLQP-21 mice depends on adaptive thermogenesis requirements

Based on the unexpected resistance to diet-induced obesity, and considering the previously established role for TLQP-21 in energy balance, we tested the hypothesis that the metabolic phenotype was caused by a compensatory response to sub-thermoneutral housing at room temperature which is known to drive sympathetic tone and norepinephrine/β-adrenergic receptor (NE/βAR) signaling to metabolic organs in order to sustain body temperature and metabolism ^36–38^. To test this hypothesis, we housed ΔTLQP-21 and WT mice at 28-30°C, a temperature selected to minimize adaptive thermogenic demands. Supporting our hypothesis, this housing condition was sufficient to prevent resistance to HFD-induced obesity in ΔTLQP-21 mice as evidenced by body weight, food intake, body composition and glucose homeostasis, all of which resulted in non-significant differences from WT mice (**Figure 6**). In addition, no significant differences were found for energy expenditure, fat-free mass, and food intake (**Figure 6; Supplementary Figure 7**). Interestingly, while some of the thermogenic genes were normalized by these housing conditions, *Ucp1* and *Cox71a* were still significantly elevated in ΔTLQP-21 mice (**Figure 6I**).

**Figure 6.**
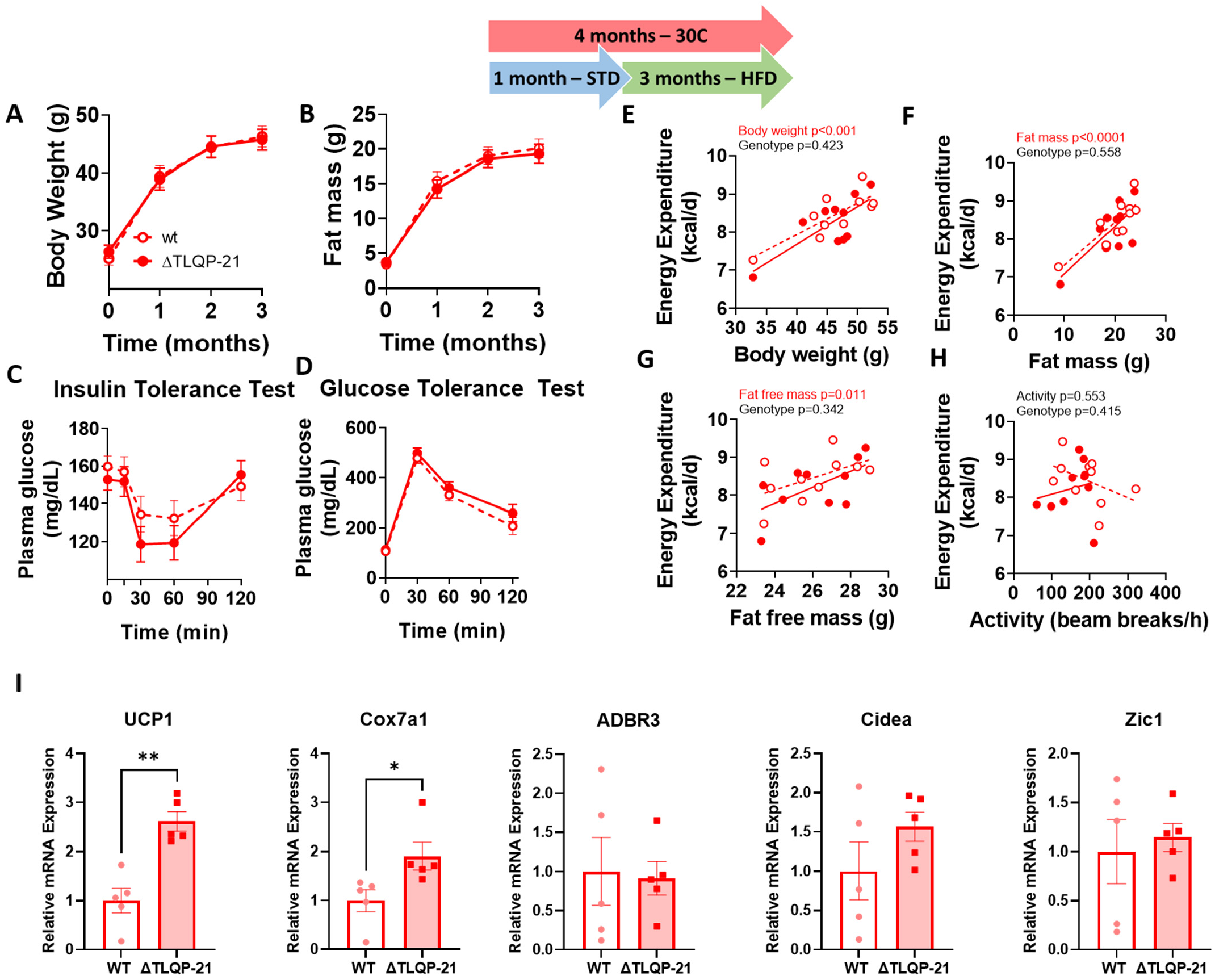
Metabolic phenotypes of male ΔTLQP-21 mutant and wild type mice tested at 30±1°C and ad libitum fed a 60% High Fat Diet (HFD). ΔTLQP-21 mice did not differ from WT mice in either body weight or fat mass gain over time (A,B), nor in glucose homeostasis tests (C,D). Energy expenditure assessed during the course of 24h showed no dependency on genotype but a significant dependency on mouse body weight (E), fat mass (F), and fat free mass (G), with no relationship to activity levels (H). N=10/group. (I) QPCR of genes implicated in brown adipocyte thermogenesis and browning. UCP1 (p=0.0010), Cox7a1 (p=0.0343). Data were analyzed using two-tailed unpaired t-test. Symbols represent individual ΔΔCt values and normalized to WT. * represents p<0.05, ** represents p<0.01.

## Discussion

Here we developed and initially characterized a mouse model carrying the selective loss of the VGF-derived TLQP-21 peptide leaving expression of the VGF pro-protein intact. Additionally, we showed that preventing the cleavage of TLQP-21 does not significantly increase the accumulation of its direct precursor TLQP-62 ^16^, a result that can be ascribed to the rapid proteolytic cleavage of small peptides ^24,29^. The generation of the ΔTLQP-21 mouse was achieved by generating a knock-in mouse carrying a selective point mutation in the codon encoding for R_21_ that is required for its proteolytic cleavage (and is also required for its biological activity via C3aR1 activation ^1,16,17^). Validation of this model included: i) sanger sequencing of the gene; ii) a novel in-gel digestion targeted proteomics which allowed unambiguous confirmation of the model by identifying a non-natural fragment (HFHHALPPA**<u>A</u>**HHPDLEAQAR) created by the R_21_→A mutation; iii) absence of immunoreactivity to a selective anti-TLQP-21 antibody ^20,30^.

Despite previous work suggesting a role for TLQP-21 in sexual behavior and reproduction ^39–41^, ΔTLQP-21 mice reproduce well and perform normal maternal behavior. Additionally, when mice are fed a standard diet, the R_21_→A mutation does not confer gross behavioral or metabolic abnormalities except for a defective acute cold-induced lipolysis, which is consistent with a role for TLQP-21 to potentiate adrenergic induced lipolysis *in vitro* and *in vivo* ^20,23,24,33^.

Based on the anti-obesity effect exerted by central/peripheral TLQP-21 injection in mice or hamsters ^16,33,34^, we initially hypothesized that ΔTLQP-21 mice would be vulnerable to HFD-induced obesity. Contrarily to our hypothesis, ΔTLQP-21 mice were resistant to HFD-induced obesity, a phenotype that cannot be explained by changes in food intake, activity or, surprisingly, energy expenditure, which were all unaffected by the ablation of TLQP-21. It is possible that technical limitations in the use of indirect calorimetry ^42,43^ prevented the detection of increased energy expenditure and/or that energy compensation mechanisms were activated ^44–46^. Two lines of evidence suggest this conclusion. Firstly, the BAT of ΔTLQP-21 mice showed increased markers of activation when compared to WT, suggesting increased thermogenic capacity. Secondly, minimizing adaptive thermogenesis requirements by housing mice at 28-30°C prevented the obesity resistant phenotype from manifesting. It is thus possible that selective germline deletion of TLQP-21 caused a compensatory augmentation of other thermogenic pathways overall resulting in a paradoxical resistance to obesity despite a catabolic role for the peptide itself. While this hypothesis requires additional experimental support to be mechanistically understood, two lines of evidence seems to indirectly support it. First, we detailed the existence of a strong selective pressure in mammals on the C-terminus of TLQP-21 (see ^23^ and **Figure 1**). We showed that the S_20_→A mutation is exclusive to *Murinae* and *Cricetinae* and conferred a gain of function resulting in enhanced potency at the C3aR1 receptor to regulate lipolysis ^23^. We speculate that selective pressures favored the emergence of a pro-lipolytic and thermogenic mechanism in small-sized rodents. In the absence of this mechanism, due to genetic loss of function, one of two outcomes would be possible: i) ablating the peptide could result in hypothermia and/or defective lipolysis and obesity; ii) due to its physiological relevance, the genome/phenome could respond by activating and/or increasing the efficacy of other catabolic pathways. That the second scenario can be correct is suggested by the consistent (thus far considered paradoxical) resistance to obesity manifested by other models including VGF-/-^11,12^, the C3aR-/-^35^ as well as a humanized mouse carrying a selective truncation of a large portion of the VGF C-terminus including TLQP-21 ^15^. It is thus possible that lack of TLQP-21 may be the main reason why VGF-/-mice were resilient to HFD-induced obesity ^11,12^. Furthermore, a humanized knock in mouse carrying a full length human VGF, including the S_20_ at the C-terminus of TLQP-21, is prone to obesity ^15^. Overall, these results suggesting that the ancestral -SR_21_ sequence ^23^ which is shared with humans can cause the pro-obesogenic phenotype in mice which was predicated based on the catabolic effect of pharmacological treatment of WT mice/adipocytes with TLQP-21 ^20,33^. Conversely, the absence of TLQP-21 in the germline of ΔTLQP-21 and in the VGF-/-mouse models, could trigger the activation of compensatory physiological mechanisms leading to the obesity resistant phenotype. Lastly, we cannot rule out a developmental role for the TLQP-21 peptide in metabolic tissues, such that TLQP-21 ablation in the germline (of either the ΔTLQP-21 or the VGF-/-^11^) results in altered BAT morphology and activity, and resistance to HFD-induced obesity, while peptide administration in the adult animal generally has the opposite effect.

### Conclusion and future perspectives

Here we developed a mouse model of selective loss of TLQP-21. The ΔTLQP-21 line can be a valuable resource to conduct mechanistic studies on the necessary role of TLQP-21 in physiology and disease, while also serving as a platform to test the specificity of novel antibodies or immunoassays directed at TLQP-21. We focused our characterization on energy balance, metabolism and behavior, which revealed a unique role for this peptide in energy balance in response to dietary and environmental challenges. We did not test other biological roles attributed to TLQP-21 such as in nociception, gastrointestinal functions, blood pressure, and others. Thus, additional work is necessary to characterize the physiology of this mouse model in full.

Aside from testing specific hypotheses relevant to VGF and TLQP-21 biology, our approach can have far-reaching implications by informing the development of similar knowledge-based genetic engineering approaches to generate selective loss of function of other peptides encoded by pro-hormones genes, leaving all other peptides within the pro-protein precursor intact and unmodified. These studies can open new vistas to further characterize the biology of many biologically active peptide hormones encoded by pro-peptides whose function has been studied thus far using mostly gain of function or complete loss of function gene ablation approaches.

## Methods

### Generation of Vgf ΔTLQP21 mouse line

The floxed VGF ΔTLQP21 (R_21_→A) knockin mouse line was generated by inserting a 5’ flanking loxP site into the Vgf 5’ UTR (KpnI site), and a 3’ flanking loxP site and FRT-flanked neomycin selection cassette, derived from p-loxP-2FRT-PGKneo (Dr. David Gordon, University of Colorado Health Science Center), into the Vgf 3’ UTR (XbaI site), using previously described mouse Vgf genomic sequences ^11^. This previously described VGF construct ^27^ underwent targeted mutagenesis (Mutagenex Inc., Hillsborough NJ) to introduce a mutation encoding a TLQP-R21A amino acid substitution (mVGF_1-617_ R576A). The plasmid sequence was verified and the targeting construct was electroporated into 129Sv/J-derived R1 embryonic stem (ES) cells by the Mouse Genetics and Gene Targeting Core Facility, Icahn School of Medicine at Mount Sinai, as described previously ^11,47^. Southern and PCR analyses confirmed correct gene targeting. Male chimeras were mated with C57BL/6J females to produce F1 breeders and experiments were performed on N2F1 mice. Western blot analysis of brain lysates using anti-VGF (C-term) demonstrated that the mVGF_1-617_ R576A protein (containing TLQP-R21A) was the anticipated size and was expressed from the targeted allele in comparable amounts to wild type mVGF_1-617_.

### Genotyping and Sanger sequencing

Genomic DNA was isolated from the mouse tails using the Qiagen DNA extraction kit. The VGF region of interest was PCR amplified with specific primers VGF-P.11-for 5’-TCACCCCTTCCCAAACTACA-3’ and VGF-P.11-rev 5’-ATTCTCCAGCTCCTCCTGCTC-3’, and the resultant amplicon was gel purified. Gel-purified amplicons were subjected to Sanger sequencing by using the same primers. The results were analyzed using Chromas Lite software, and base changes in the ΔTLQP21 mice were confirmed by manually inspecting the peaks of the chromatogram. The TLQP-21 region of the Vgf gene was PCR amplified and sequenced at the University of Minnesota Genomics Center. Chromatograms were manually analyzed using Chromas Lite software for the mutations, and the peaks were extracted for the same.

### Mass spectrometry

For mass spectrometry, fresh tissues (3 adrenal glands) for WT and ΔTLQP21 were collected and lysed in lysis buffer (2.5% SDS, 100mM Tris, pH 8) using a bead-based homogenizer. Lysates were cleared; the BCA method was used to estimate protein (Thermo Fisher). Equal amounts of protein were loaded in each well and resolved on 12% gels (Invitrogen, Bolt) with MES buffer. A molecular weight marker ladder (Dual Extra Plus, Bio-Rad) was used. In parallel, the synthetic peptide was also loaded for precise gel excision of the TLQP-21 region. Gels were resolved at 200V for 30 minutes, washed with HPLC grade water and stained using safe blue stain (Invitrogen) for 60 minutes, followed by de-staining using HPLC grade water. With the synthetic peptide and the molecular weight marker as a reference, gel bands were excised using sterile scalpels and stored at -20 degrees until further processing. In-gel digestion was carried out per standard protocols, and the tryptic peptides were used for further analysis. Peptide MS Analysis: peptide digestions were dried and solubilized in 20ul of 2% acetonitrile, 98% water and 0.1% formic acid. 2ul was loaded on an Eksigent 2.7uM HALO fused-core c18 column (100mm x 0.3mm) using an Agilent 1100 series microflow pump. The peptides were loaded on the column at a rate of 8ul/min for 2 minutes and a linear gradient from 2% acetonitrile (ACN) to 45% ACN for 17.5 minutes. The ACN was then increased to 90% for 2 minutes and then to 2% ACN for 5 minutes. The Applied Biosystems 5500 ion trap was used using a turbo V electrospray source fitted with a 25uM ESI electrode. Transitions and parameters predicted from an insilico tryptic digestion of the synthetic TLQP-21 peptide with all partially digested fragments considered. The tryptic-digested synthetic peptide validated the MS parameters determined using the Skyline program ^48^. The data was analyzed using MultiQuant™ (ABI Sciex, Framingham, MA, USA), which provides the peak areas for the transitions.

### General in vivo procedures

Experimental mice were maintained in a fully controlled animal facility (12:12 h light:dark cycle at 22 ± 2°C, unless otherwise specified). Heterozygous breeding pairs were established and pups were weaned in groups of same-sex siblings. Animal experiments were conducted at the University of Minnesota (USA) and approved by the Institutional Animal Care and Use Committee, University of Minnesota. Mice were fed a standard (D12450B, Research Diets 3.85 kcal/g, 10% kcal from fat) or a high fat (D12492, Research Diets, 5.24kcal/g, 60% kcal from fat, HFD) diet.

Mice undergoing experiments at thermoneutrality, were transferred from the vivarium to an adjacent fully controlled environmental room and housed in a 12:12 h light:dark cycle at 30± 1°C. Mice were allowed one month to acclimate to 30± 1°C before performing any experimental procedure.

In separate studies, ΔTLQP-21 and WT male and female mice (n=8-10/group) were single housed and assessed for: body weight, food intake, body composition, glucose homeostasis, energy expenditure and behavior, cold and β-adrenergic receptor sensitivity. The study design also incorporated environmental factors such as housing temperature (standard housing temperature 22°C or mouse thermoneutral temperature 28-30°C) and diet (standard diet or high fat diet). Body weight and food intake were measured regularly during the experiments.

### Maternal behavior

Maternal behavior was assessed by observing ΔTLQP-21 (n=6), WT (n=5) and heterozygous (n=7) mothers in their home cages, from post-natal day (PND) 1 to 14. Dams behavior was analyzed in the last 2h of the light cycle by the same operator who was unaware of the mouse genotype. Each lactating female was observed once every 4 minutes, for a total amount of 30 observations in 2 hours for each dam. During each 4min observation, the experimenter recorded the behavior displayed at the moment of the observation. Dams behaviors were identified according to standard behavioral categories ^49^ including: arched back (AB) nursing, nursing, licking, nest building, grooming, active. We also calculated the time spent out of the nest; considering all the observations during which the dam was anywhere in the cage but the nest, regardless of the behavior exhibited at the moment of the observation. In addiction we put together all behaviors related to pups (AB, nursing, nest building, licking pups) and we compared it with total of behaviors not related to pups (out of nest, active, grooming).

### 4h fasting glucose, Glucose Tolerance Test, Insulin Tolerance Test

Fasting glucose was measured 4h following food removal by tail bleeding. Glucose tolerance test was performed following an overnight fast. Blood glucose levels from tail bleeding were monitored at 0, 30 60 and 120 min after an i.p. injection of D-glucose at 2 g/Kg. Insulin tolerance test was performed following a 6 h fasting period. Mice were injected i.p. with a dose of 0.75 IU/kg of insulin and blood glucose levels were monitored at 0, 15, 30, 60, and 120 min after injection. Glucose was measured with Acu-check Aviva glucometers (Roche Diagnostics, Indianapolis, IN).

### Indirect Calorimetry and Body Composition

Oxygen consumption (VO_2_), carbon dioxide production (VCO_2_), Heat, Respiratory Efficiency Ratio (RER) and activity were measured using the Oxymax Comprehensive Lab Animal Monitoring System (Columbus Instruments). Energy expenditure was calculated with the formula provided by the manufacturer, expressed as kcal/h and analyzed with body weight, fat mass, fat free mass or activity as continuous predictor in an ANCOVA model. Body composition was measured with EchoMRI 3-in-1 (EchoMRI LLC, Houston, TX, USA). Variables were measured in a rapid (approximately 70 seconds) and noninvasive manner and included total body fat and fat free mass.

### Elevated Plus Maze Test

The Elevated Plus Maze (EPM) for mice consisted of intersecting perpendicular open arms (60 × 5 cm) and enclosed arms (60 × 5 cm) with 20 cm high walls. The maze was raised to a height of 50 cm above the floor level in a dimly lit room (20 Lux) and a video camera was suspended above the maze to record the movements for analysis. Each mouse was placed at the center of the platform, its head facing an open arm. The animals were tested individually and only once for 5 min. The maze was cleaned after each trial so as to remove any residue or odors. The following measurements were automatically taken and analyzed using a video-tracking software (Ethovision XT, Noldus, The Netherlands): the number of entries into the open or closed arms, the time spent in each arm, and the total distance moved in the EPM.

### Forced Swim Test

Mice were individually forced to swim in an open cylindrical glass container (diameter 10 cm, height 25 cm), containing 10 cm of water at 25 ± 1°C, for 6 min. The water was changed before the introduction of each animal. Mouse behavior was video-recorded by a video-camera placed in front of the glass cylinders. Activity was measured from the video recordings using video tracking software (Ethovision XT, Noldus, The Netherlands). Activity was scored using the mobility threshold settings within the Ethovision software by measuring the percentage change in area of the tracked object from one 0.08 sec time bin to the next. Activity was defined as either immobile (for changes in area between 0–10%) or strong mobile (for changes in area between 60–100%) during the last 4 min of the test.

### Glycerol assay during cold and β3-AR challenge

To evaluate the acute thermogenic responses to cold, mice were moved to a fully environmentally controlled environment where the thermostat was set at 4°C. For the selective β3 adrenergic receptor agonist challenge, CL-316243 (Sigma-Aldrich) dissolved in sterile saline was administered i.p. at 1mg/kg. For both procedures, mice were sampled 3h after either the exposure to cold or the CL-316243 injection.

Glycerol was measured in mouse plasma samples using the Glycerol Assay Kit (MAK117, Sigma-Aldrich) following the procedure recommended by the manufacturer. Glycerol standards were diluted with water to get final concentrations of 1.0, 0.6, 0.3, and 0mM. Plasma samples were diluted with water, using a dilution factor of 4. The colorimetric assay was conducted in a 96-well plate and absorbance was measured using a plate reader (BioTek) at 570nm. Using the glycerol standards, a standard curve was plotted and the slope was determined using linear regression. The measured absorbance values of the sample were first multiplied by the dilution factor, and then plasma glycerol concentrations were calculated using the measured absorbance of the sample divided by the slope of the standard curve.

### Insulin ELISA

Plasma insulin levels was measured using Mouse Ultrasensitive Insulin ELISA (ALPCO) according to kit instructions.

### Immunohistochemistry in frozen tissues

Tissue from wild-type or ΔTLQP-21 mutant animals was harvested and fixed overnight in 4% PFA/PBS at 4°C. The tissue was then dehydrated by storage in 30% (wt/vol) sucrose for a minimum of two days. Adrenal tissue was embedded in OCT (optimal cutting temperature) compound using dry ice-cooled ethanol and cryosectioned (−20°C) at a thickness of 20μm mounted directly to charged slides (Fisherbrand Superfrost Plus). Slides containing collected cryosections were stored at -80°C for later use. Collected adrenal sections were probed with antibodies for TLQP-21 (rabbit anti-TLQP-21, 1:100, ^20^) or Tyrosine Hydroxylase (sheep polyclonal TH, 1:100, Millipore Sigma) overnight at 4°C in a humidified chamber. The sections were the labeled with Alexa Flour-conjugated secondary antibodies (donkey anti-rabbit 488, donkey anti-sheep 647, 1:500; Jackson ImmunoResearch Laboratories), counterstained with DAPI, and coverslips mounted using ProLong Gold mounting media (Invitrogen).

### Morphological analysis of Formalin-Fixed Paraffin-Embedded (FFPE) adipose fat pads

Immediately after removal, tissue pieces representing BAT, ScWAT and pWAT were fixed in 4% paraformaldehyde in 0.1 M phosphate buffer (pH 7.4) by overnight immersion at 4°C. The samples were then dehydrated, cleared, and embedded in paraffin. Serial sections, from 3 different levels (100 μm apart), were respectively H&E-stained to assess their morphology and immunostained for UCP1. Immunostaining was performed as follows: 3-μm-thick sections were dewaxed, incubated with rabbit anti-UCP1 (1:500; ab10983, Abcam, Cambridge UK) according to the avidin-biotin complex. Briefly: 1) endogenous peroxidase blocking with 3% hydrogen peroxide in methanol; 2) normal serum blocking (1:75) for 20 min to reduce nonspecific background; 3) incubation with primary antibodies against UCP1 at 4°C; 4) secondary antibodies specific for rabbit, IgG biotin conjugated (1:200; Vector Labs, Burlingame, CA, USA); 5) ABC kit (Vector Labs); and 6) enzymatic reaction to reveal peroxidase with Sigma Fast 3,3’-diaminobenzidine (Merck KGaA, Darmstadt, Germany) used as substrate. Finally, sections were counterstained and mounted in Eukitt (Fluka, Deisenhofen, Germany). All observations were performed with a Nikon Eclipse 80i light microscope (Nikon, Tokyo, Japan) equipped with a CCD camera. Brightness and contrast of the final images were adjusted using the Photoshop CS3 software (Adobe Systems; Mountain View, CA, USA).

### qPCR

Total RNA was isolated from frozen tissue following the protocol listed in the RNeasy Lipid Tissue Mini Kit by Qiagen (Catalog #74804). cDNA synthesis from RNA was completed using 5x iScript RT Supermix (Bio-Rad, Catalog #L001404 B) according to the manufacturer’s protocol. Individual cDNA samples were run in duplicate using iTaq Universal SYBR Green Supermix (Bio-Rad, Catalog #L001752 B), with a final volume in each well equal to 15uL. A CFX Connect Real-Time System (Bio-Rad) thermal cycler and optic monitor was used. Gene expression data was normalized to the geometric mean of the best housekeeping gene, GAPDH, as determined by the program RefFinder ^50^. Data was analyzed using the ΔΔC_t_ method and further normalized to the control group’s mean. Primers are listed in **Supplementary Table 5**.

### Statistical analysis

Statistical significance was determined by one or two way ANOVA or ANOVA for repeated measures with Tukey’s post hoc test, with Student’s t-test or with the non-parametric Mann-Whitney U test where appropriate. Statistica (TIBCO Software Inc.) was used for analyses.

## Acknowledgement

Kohinoor Khan, Vinayak Ghosh are acknowledged for help with the study execution and figure preparation respectively.

