## Supplementary Figures for "Targeted and selective knockout of the TLQP-21 neuropeptide unmasks its unique role in energy homeostasis"

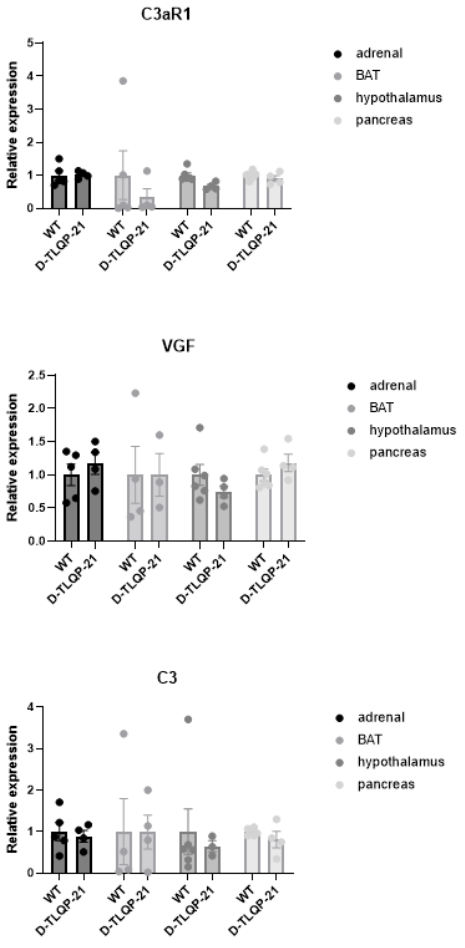


**Supplementary Figure 1. Expression of VGF, C3 and C3aR1 in various organs of male mice fed a standard diet.** No statistical difference was detected between WT and ∆TLQP-21 mice. N=4-5.

**WT**

**Δ-TLQP-21**

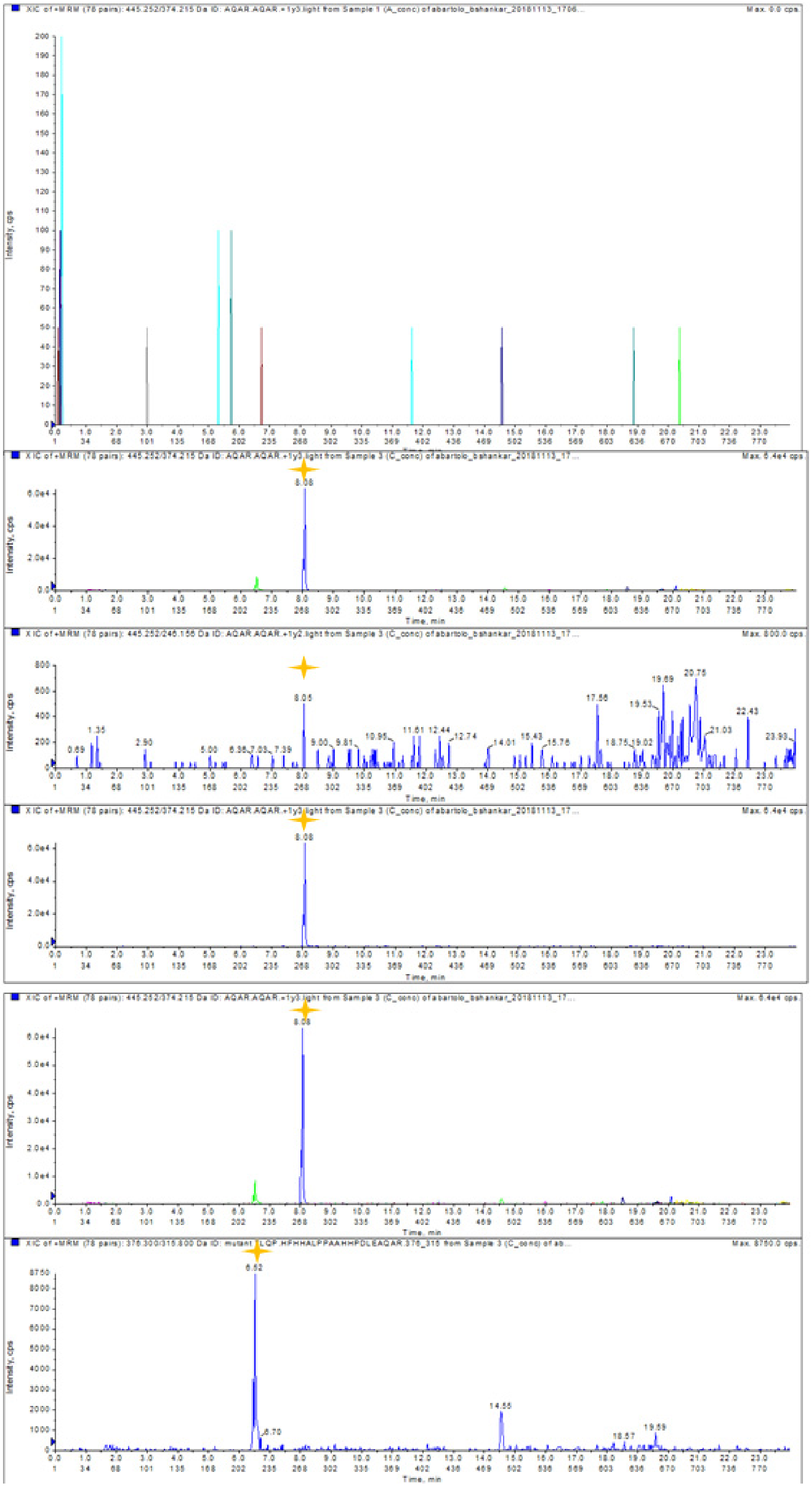


**Supplementary Figure 2.** MS Raw data: Representative mass spec data for the unnatural TLQP-21 fragments, detected only in delta TLQP-21 mice (marked by a star) but not in wild type mice as predicted by the targeted mass spectrometry software Skyline.

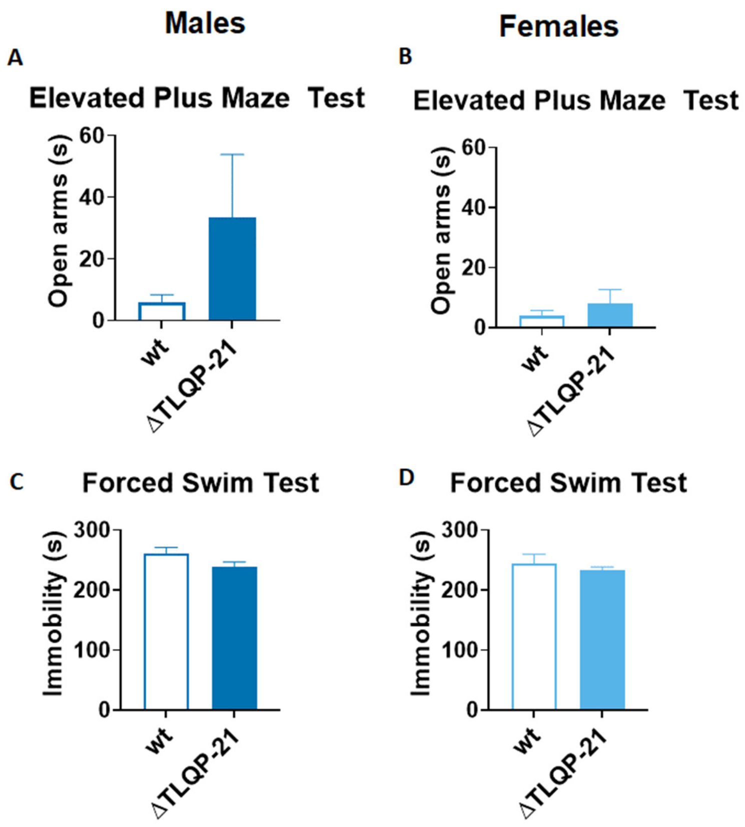


**Supplementary Figure 3. Behavioral characterization of both males and females ΔTLQP-21 mutant and WT mice**. No difference in anxiety-like behaviors assessed in the elevated plus maze test (**A,B**) and no differences in depression-like behaviors assessed in the forced swim test (**C,D**) were observed. N=8-10/group.

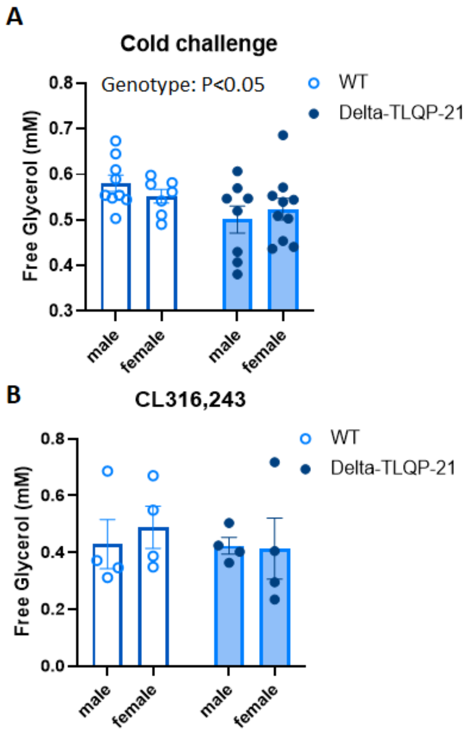


**Supplementary Figure 4. Lipolysis in response to cold or β3 Adrenergic Receptor (AR) challenge. A)** Without any sex difference, ΔTLQP-21 mice show a lower response to 3h exposure to 4°C when compared to WT ($F(1,31)=5.49$, $p=0.0257$). **B)** No genotype or sex difference is detected in lipolysis 3h following injection of the β3AR agonist CL316,243 (1 mg/Kg).

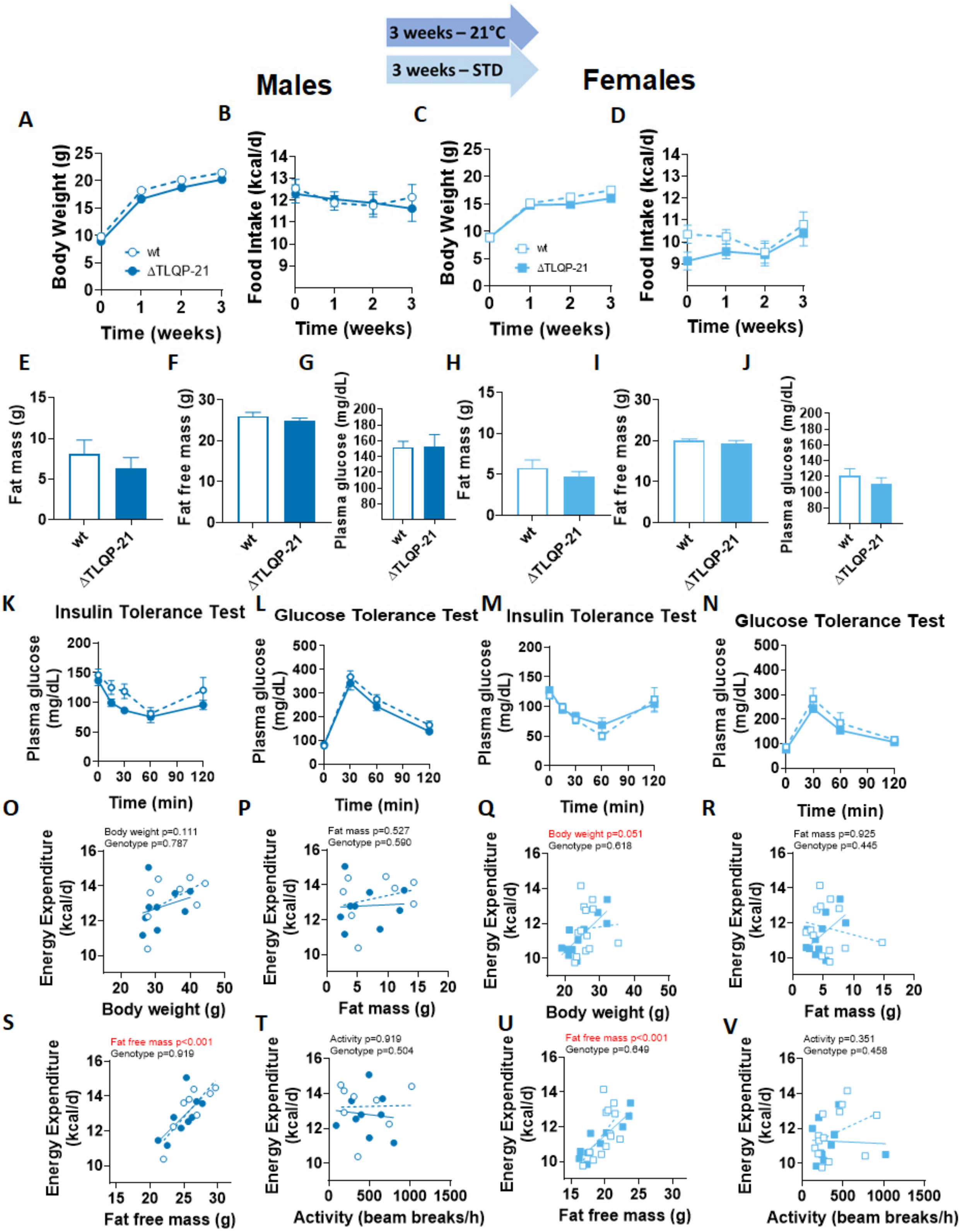


**Supplementary figure 5. Metabolic characterization of male and female ΔTLQP-21 mutant and WT mice tested at 21±2°C and fed ad libitum standard diet.** In both males and females, ΔTLQP-21 and WT mice had similar body weight gain (**A,C**), food intake (**B,D**), body composition (**E-F, H-I**), plasma glucose levels after 4h fasting (**G,J**), and glucose homeostasis (**K-L, M-N**). Energy expenditure assessed over 24h, showed no significant dependency on genotype, while being significantly influenced by body weight and fat free mass in females (**Q**), and fat free mass in males (**S**). N=8-13/group.

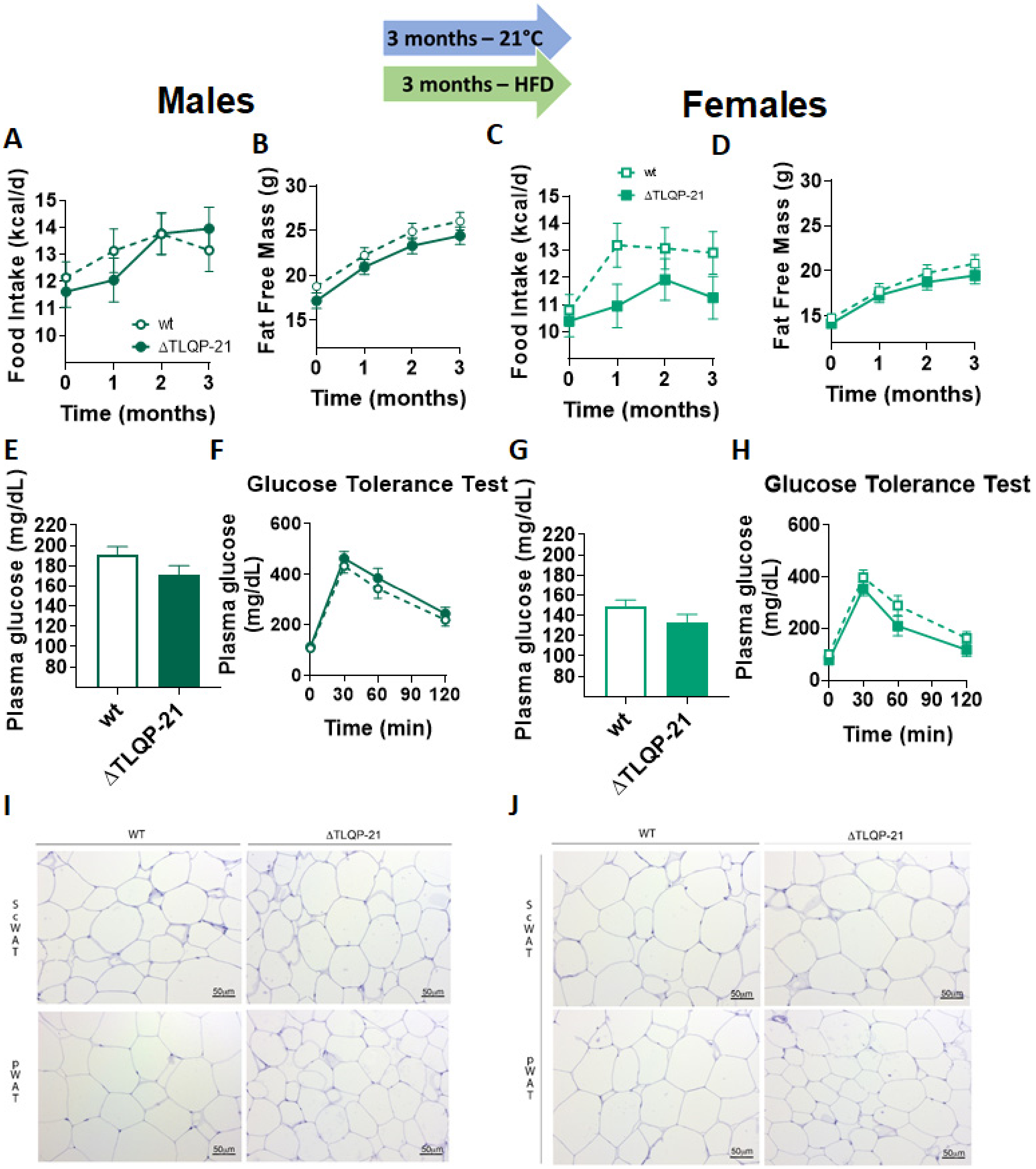


**Supplementary figure 6. Metabolic characterization of ΔTLQP-21 and WT male and female mice tested at 21±2°C and ad libitum fed 60% High Fat Diet (HFD).** In both males and females, there were no differences between ΔTLQP-21 and WT mice in food intake **(A,C)**, fat free mass **(B,D)**, plasma glucose levels after 4h fasting **(E, G),** and glucose tolerance **(F,H)**. N=6-9/group. **I,J**) Morphological analysis show that the pWAT but not scWAT of male mice fed HFD had a higher proportion of smaller size adipocytes in the ΔTLQP-21 mice (N=4).

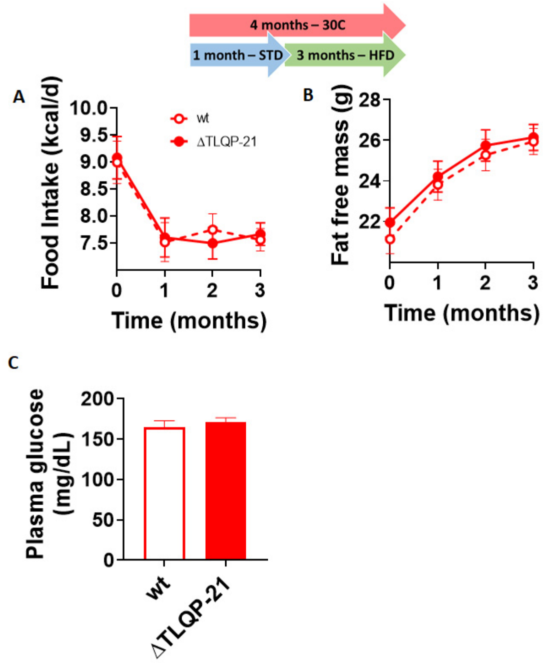


**Supplementary figure 7. Metabolic characterization of ΔTLQP-21 mutant and WT male mice tested at 30±1°C and ad libitum fed 60% High Fat Diet (HFD).** No differences emerged in in food intake (**A**), fat free mass (**B**), and plasma glucose levels after 4h fasting (**C**). N=10/group.
