## Supplementary Table 2 for "Targeted and selective knockout of the TLQP-21 neuropeptide unmasks its unique role in energy homeostasis"

**Supplementary Table 2. Reproductive success.**

| Breeder type | # pups | # males | # females | Sex ratio |
| --- | --- | --- | --- | --- |
| Heterozygous (n=19) | 6.21 ± 0.6 | 3.58 ± 0.45 | 2.63 ± 0.39 | 0.53 ± 0.06 |
| Homozygous (n=40) | 6.60 ± 0.42 | 3.5 ± 0.31 | 3.1 ± 0.27 | 0.53 ± 0.14 |
| WT  (n=24) | 7.21 ± 0.53 | 3.71 ± 0.40 | 3.5 ± 0.34 | 0.51 ± 0.05 |
| ΔTLQP-21  (n=16) | 5.69 ± 0.65 | 3.18 ± 0.49 | 2.5 ± 0.41 | 0.56 ± 0.06 |

The number of pups (total, males and females) is expressed per litter

Sex ratio is calculated as “number of males per litter/(total number of pups per litter)”.
