## Supplementary Table 3 for "Targeted and selective knockout of the TLQP-21 neuropeptide unmasks its unique role in energy homeostasis"

|  | Breeders | | | |
| --- | --- | --- | --- | --- |
| **%Behavior** | Heterozygous (n=5) | Homozygous  (n=9) | WT  (n=4) | ΔTLQP-21  (n=5) |
| Arched back nursing | 59.81 ± 3.14 | 56.72 ± 2.34 | 60.24 ± 2.92 | 53.90 ± 3.52 |
| Nursing | 8.76 ± 1.26 | 8.26 ± 1.26 | 4.64 ± 1.37 | 11.14 ± 1.53 |
| Pup licking | 6.86 ±0.96 | 4.60 ± 0.71 | 5.00 ± 0.90 | 4.28 ± 0.34 |
| Nest building | 3.04 ± 0.50 | 3.07 ± 0.38 | 3.33 ± .019 | 2.86 ± 0.54 |
| Self-grooming | 6.95 ± 1.40 | 8.62 ± 1.04 | 10.71 ± 2.13 | 6.95 ± 1.15 |
| Out of nest | 17.43 ± 3.28 | 21.32 ± 2.44 | 19.40 ± 3.03 | 22.86 ± 4.60 |
| Active | 21.62 ± 3.33 | 24.13 ± 2.48 | 21.07 ± 4.07 | 26.557 ± 3.70 |
| Total pup related | 78.47 ± 3.92 | 72.64 ± 2.92 | 73.21 ± 4.29 | 72.19 ± 4.36 |
| Total pup unrelated | 46.00 ± 6.03 | 54.07 ± 4.49 | 51.19 ± 6.04 | 55.38 ± 8.11 |

**Supplementary Table 3. Maternal behavior**

Behaviors are expressed as weekly percentages.
