## Supplementary Table 4 for "Targeted and selective knockout of the TLQP-21 neuropeptide unmasks its unique role in energy homeostasis"

**Supplementary Table 4. Body weight at weaning**

| Gender | Genotype | Breeder | Rel. Body Weight at Weaning | N |
| --- | --- | --- | --- | --- |
| Male | WT | Heterozygous | 2.05 ± 0.33 | 9 |
|  |  | Homozygous | 1.70 ± 0.27 | 18 |
|  | ΔTLQP-21 | Heterozygous | 1.61 ± 0.16 | 10 |
|  |  | Homozygous | 1.58 ± 0.24 | 4 |
| Female | WT | Heterozygous | 1.95 ± 0.27 | 9 |
|  |  | Homozygous | 1.22 ± 0.16 | 8 |
|  | ΔTLQP-21 | Heterozygous | 1.50 ± 0.14 | 9 |
|  |  | Homozygous | 1.72 ± 0.16 | 10 |

Since litters weren’t culled at birth, the presented body weight is the individual body weight measured at weaning divided by the litter size.
