## Supplementary Table 5 for "Targeted and selective knockout of the TLQP-21 neuropeptide unmasks its unique role in energy homeostasis"

| Primer | Forward Sequence (5’ to 3’) | Reverse Sequence (5’ to 3’) |
| --- | --- | --- |
| ADBR1 | CTC ATC GTG GTG GGT AAC GTG | ACA CAC AGC ACA TCT ACC GAA |
| ADBR2 | GGG AAC GAC AGC GAC TTC TT | GCC AGG ACG ATA ACC GAC AT |
| ADBR3 | CCA GCC AGC CCT GTT G | GGA CGC GCA CCT TCA TAG C |
| B-Actin | GGC ACC ACA CCT TCT ACA ATG | GGG GTG TTG AAG GTC TCA AAC |
| Cidea | TGC TCT TCT GTA TCG CCC AGT | GCC GTG TTA AGG AAT CTG CTG |
| Cox7a1 | CAG CGT CAT GGT CAG TCT GT | AGA AAA CCG TGT GGC AGA GA |
| C3aR1 | TCG ATG CTG ACA CCA ATT CAA | TCC CAA TAG ACA AGT GAG ACC AA |
| GAPDH | AGG TCG GTG TGA ACG GAT TTG | TGT AGA CCA TGT AGT TGA GGT CA |
| GPR3 | ATC ACC TGA GCA ACC GAG AA | AGA TGG GGG TGC ATT TTA CA |
| P2RX5 | CAC GAT GCT TGG GGA ACG ACT | CCG CTG GTT AGG AGT CAC GAT |
| UCP1 | GTC CCC TGC CAT TTA CTG TCA G | TTT ATT CGT GGT CTC CCA GCA TAG |
| Zic1 | AAC CTC AAG ATC CAC AAA AGG A | CCT CGA ACT CGC ACT TGA A |
